# Nonreciprocal synchronization in embryonic oscillator ensembles

**DOI:** 10.1101/2024.01.29.577856

**Authors:** Christine Ho, Laurent Jutras-Dubé, Michael Zhao, Gregor Mönke, István Z. Kiss, Paul François, Alexander Aulehla

## Abstract

Synchronization of coupled oscillators is a universal phenomenon encountered across different scales and contexts e.g., chemical wave patterns, superconductors and the unison applause we witness in concert halls. The existence of common underlying coupling rules define universality classes, revealing a fundamental sameness between seemingly distinct systems. Identifying rules of synchronization in any particular setting is hence of paramount relevance. Here, we address the coupling rules within an embryonic oscillator ensemble linked to vertebrate embryo body axis segmentation. In vertebrates, the periodic segmentation of the body axis involves synchronized signaling oscillations in cells within the presomitic mesoderm (PSM), from which somites, the pre-vertebrae, form. At the molecular level, it is known that intact Notch-signaling and cell-to-cell contact is required for synchronization between PSM cells. However, an understanding of the coupling rules is still lacking. To identify these, we develop a novel experimental assay that enables direct quantification of synchronization dynamics within mixtures of oscillating cell ensembles, for which the initial input frequency and phase distribution are known. Our results reveal a “winner-takes-it-all” synchronization outcome i.e., the emerging collective rhythm matches one of the input rhythms. Using a combination of theory and experimental validation, we develop a new coupling model, the “Rectified Kuramoto” (ReKu) model, characterized by a phase-dependent, non-reciprocal interaction in the coupling of oscillatory cells. Such non-reciprocal synchronization rules reveal fundamental similarities between embryonic oscillators and a class of collective behaviours seen in neurons and fireflies, where higher level computations are performed and linked to non-reciprocal synchronization.

---

Synchronization is a universal concept that transcends across vastly different scales and contexts, including living and non-living systems [1–4]. In biology, one striking manifestation of synchronization is found during vertebrate embryonic development, as mesodermal cells establish a collective rhythm at the level of gene activity oscillations, resulting in waves that traverse the embryo along its antero-posterior axis [5]. This oscillatory activity underlies the somite segmentation clock, a molecular oscillator system that controls the periodic formation of somites, the precursors or vertebrae [6]. The molecular-mechanistic understanding of this embryonic oscillator has advanced considerably over the years and has led to the identification of several signaling pathways and essential molecular players [7–10]. Notably, oscillatory Notch-signaling pathway activity has been identified in all vertebrate species studied, including chicken [5], mouse [11], zebrafish [12], and snake [13]. Moreover, the auto-inhibition and delayed negative feedback regulation by transcriptional repressors of the Hes protein family have been shown to be at the core of Notch-signaling oscillations [14]. Notch signaling has been demonstrated to be required for cell-to-cell coupling and synchronization between presomitic mesoderm (PSM) oscillators, in both mouse [15, 16] and zebrafish embryos [17], as well as in randomized mouse PSM cell in vitro assays [18].

In contrast to this molecular-mechanistic insight, important questions regarding the basic synchronization rules of the embryonic PSM oscillator remain. For instance, it is unclear how cells are influenced by – and refer to – their neighbors’ rhythm. Are cells accelerated or delayed by the influence of neighbors? Is communication between cells bidirectional or asymmetric? To address these fundamental questions, our goal was to reveal the rules guiding synchronization between two oscillators with a similar frequency *ω*_*A*_ *≈ ω*_*B*_, but different phases *ϕ*_*A*_ */*= *ϕ*_*B*_ (Fig. 1). From a theoretical perspective, the Kuramoto model [19] is most commonly used in the segmentation clock field [20–23] so that the two oscillators are expected to adjust their phase dynamics via a sinusoidal coupling (Fig. 1B,C):

**FIG. 1.**
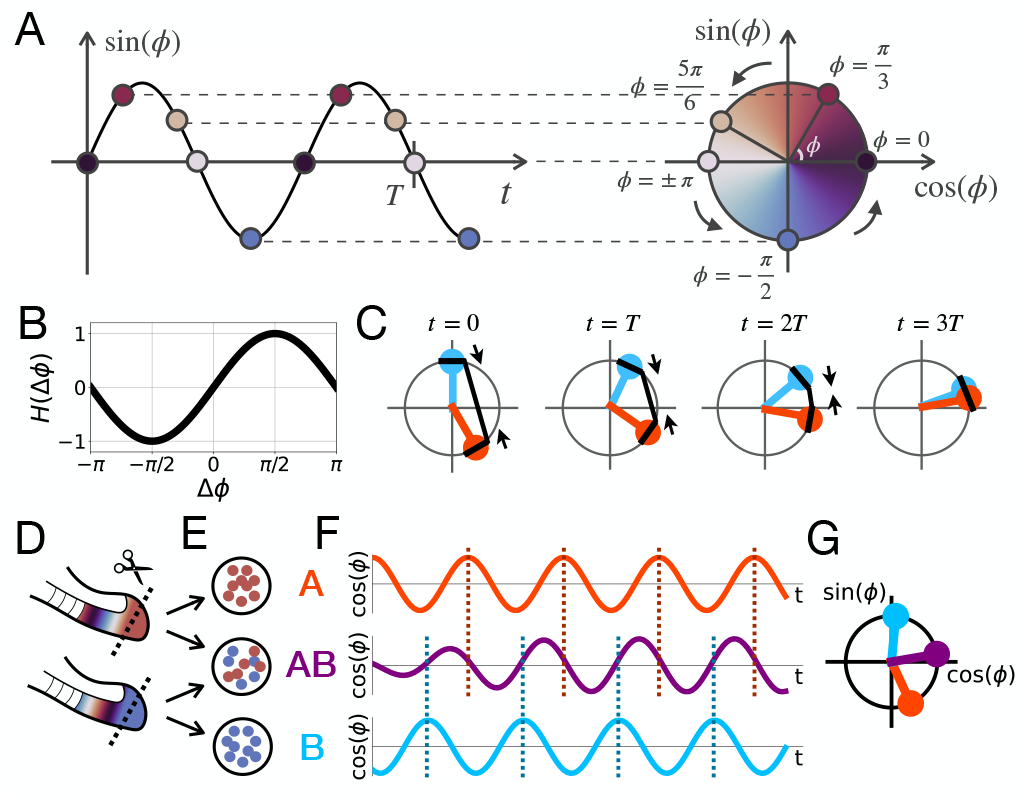
Phase synchronization dynamics. (**A**) Schematic of how an oscillating signal can be represented by a phase. (**B**) Kuramoto coupling function. (**C**) Schematic of the Kuramoto model’s phase averaging dynamics. The orange and blue dots represent *ϕ*_*A*_ and*ϕ*_*B*_, respectively. The polar plots are snapshots taken at *t* = 0, *T*, 2*T*, 3*T*, where the period *T* = 2*π/ω*, the angular frequency *ω* = *ω*_*A*_ = *ω*_*B*_ = 0.0457, and the coupling strength *c* = 0.004. The black lines and arrows schematize Kuramoto coupling, which pulls both oscillators closer. (**D** to **G**) Schematic of the experimental setup. Cells from two tail buds (**D**) are divided into three ensembles: a mixed ensemble AB, and two reference ensembles A and B (**E**). Each ensemble’s oscillations are quantified in real-time (**F**) and their phase dynamics are compared (**G**).

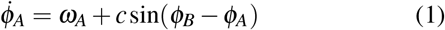

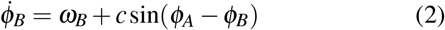

where *c* is the coupling strength (see the SI Text for more details on the origin of the Kuramoto coupling model). The outcome of Kuramoto coupling is phase averaging: the two coupled oscillators reach the average phase as they synchronize (Fig. 1F,G). Notice that the sinusoidal coupling term can take positive or negative signs depending on the phase difference *ϕ*_*A*_ *– ϕ*_*B*_, indicating a symmetric effect where both oscillators adjust their phase in response to the other oscillator, the advanced oscillator slowing down while the delayed oscillator speeds up (Fig. 1C).

While these theoretical predictions are unambiguous, their direct experimental validation remains a major challenge in biological systems, in part due to the noisy and often non-stationary oscillatory activities seen in biological oscillators. This leads to experimental difficulties to unambiguously determine, for instance, the frame of reference, i.e. the phase and frequency dynamics one would obtain without synchronization. To tackle this challenge, we thus devised a novel approach that allows just that i.e., the direct comparison of synchronization dynamics and outcome to the native, reference oscillations dynamics 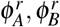. We developed a Randomization Assay For Low input (RAFL) to make a PSM cell ensemble in which cells from two different embryos are randomly mixed (population AB, phase *ϕ*_*AB*_). At the same time, we monitor a portion of the original input populations as reference (populations A and B, Fig. 1D-E), with phases 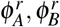. As we quantify phase and frequency behaviour using real-time imaging experiments in these three experimental conditions, i.e. A, B and AB (Fig. 1E), this novel setup enables direct quantification of the effect of synchronization, testing of theoretical predictions and more generally, identification of the underlying coupling rules.

## RESULTS

### A new experimental assay enabling randomization of single embryo oscillators

In order to obtain a cell ensemble with defined phase and frequency, we first developed an in vitro assay that tolerates low input cell numbers from a single embryo. We use only a small portion of the posterior tailbud, containing 500-1000 cells, as these share a common phase and frequency. Tailbuds are individually dissociated to a single cell suspension, split into two portions, which are then subsequently plated at high density on a fibronectin-coated coverglass, either individually (defining a reference culture) or intermixed with cells from a second tailbud. We used a fluorescent segmentation clock reporter line for Notch-signaling (LuVeLu), to quantify oscillatory dynamics within both the reference and intermixed cell ensembles, allowing for the study of synchronization (Fig. 2A-F).

**FIG. 2.**
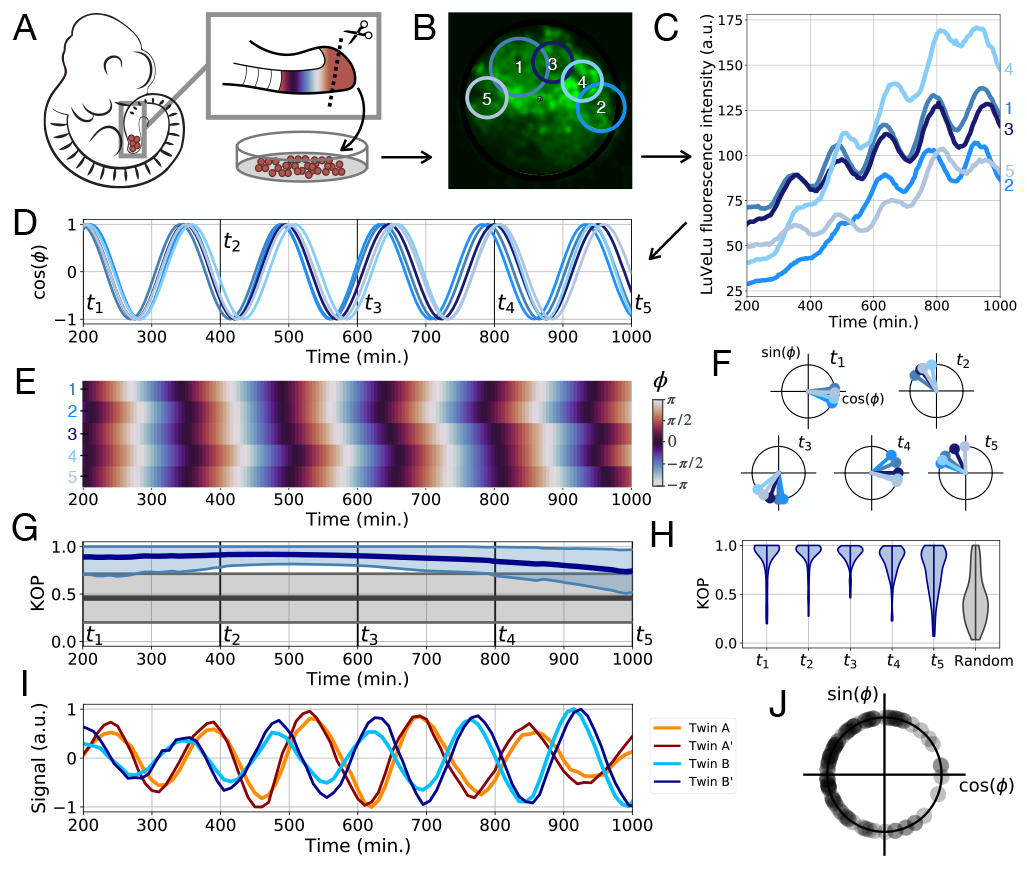
RAFL assay. (**A**) Schematic of the experimental protocol. (**B**) Oscillations are monitored in different regions of interest (ROI). (**C**) Absolute LuVeLu fluorescence in 5 ROIs. (**D**) Phase is extracted using the pyBOAT wavelet analysis toolkit [24]. (**E**) Heatmap of the phase as a function of time for the 5 ROIs. (**F**) Phase of the 5 ROIs on the unit circle every 200 minutes. Time points *t*_1_ to *t*_5_ are shown in (**D**). (**G**) Statistics of the Kuramoto order parameters (KOP). One KOP is computed for each RAFL experiment with more than one ROI (n=82). The thick dark blue line shows the mean KOP and the thin light blue lines show *±* std. The grey lines show the statistics of the KOPs of random phases with the same distribution as the experiments i.e., the total number of KOPs computed is equal to the number of experiments, and the number of random phases picked to compute each KOP is the same as the number of ROIs for the corresponding experiment. (**H**) Violin plots showing the experimental KOP distribution at each time point identified in (**G**) and the KOP distribution of random phases. (**I**) Twin experiments: two RAFLs are independently performed with cells coming from the same embryo. A and A’ show one twin experiment, and B and B’ show a second twin experiment. (**J**) Distribution of initial phases after RAFLs for all our experiments (n=85).

To validate the RAFL assay, we first quantified the synchrony within cell ensembles using the Kuramoto order parameter (KOP [20], see the SI Methods for definition). We found that within a single cell ensemble, regions of interest (ROI) at different spatial locations showed high in-phase synchrony (KOP *≈* 1) compared to randomized phases (KOP *≈* 0.5) throughout most of the time-lapse quantifications (Fig. 2, G and H). Using the circular standard deviation as an alternative for quantifying synchrony, we confirmed that each cell ensemble had a defined phase (Fig. S1). We also performed a series of validations to examine the influence of cell dissociation on the initial oscillation phase distribution. We verified that this defined synchronization outcome is reproducible by performing replicate RAFL experiments using cells from the same tailbud (Fig. 2I and Fig. S2). We also computed the phase distribution across many cell ensembles (n=85) at a given timepoint after randomization. While this distribution is slightly narrower than the distribution found in intact PSM tissue (see Fig. S3 for a comparison between tail phases and RAFL phases), the phases obtained with different RAFL experiments did range from *–π* to *π*, and hence almost spanned the full circle (Fig. 2J).

Combined, these validations showed that within each RAFL assay, the oscillation phase can be robustly determined, while at the same time demonstrating a broad phase distribution across different RAFL assays from different embryos.

### Intermixed cell ensembles from two embryos undergo winner-takes-it-all synchronization

After establishing and characterizing the RAFL experimental assay, we used it to determine the common phase upon synchronization of intermixed cells from two different embryos. More precisely, we quantified the outcome at the level of the collective phase *ϕ*_*AB*_ in the mixed population AB, and compared it to the two unmixed “reference” input population phases 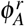 and 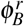 (Fig. 3A).

**FIG. 3.**
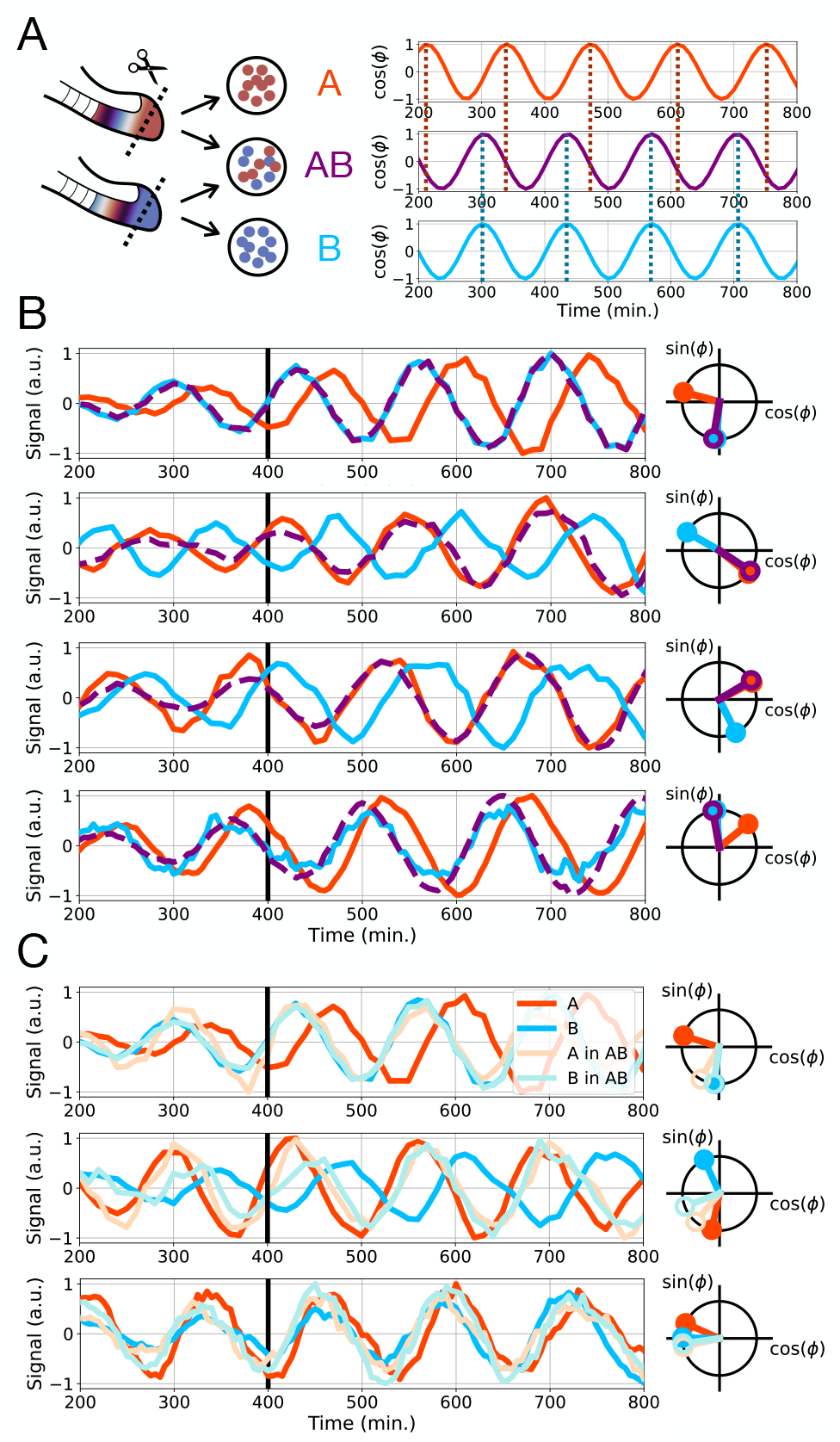
Winner-takes-it-all synchronization dynamics. (**A**) Schematic of the experimental randomization procedure and oscillation time series extracted from each cell ensemble (reference ensembles A and B and mixed ensemble AB). (**B**) Time series and polar plots of four synchronization experiments. The first row shows the data used to extract the oscillation phase time series depicted in (**A**). The thick black line indicates the time corresponding to the polar plot on the right. All experimental results (n=32) are shown in Fig. S4. (**C**) Dual-reporter imaging experiments to quantify dynamics of cells from each embryo within the mixed ensemble i.e., population “A in AB” shows the dynamics of cells originating from embryo A within the mixed population AB. The comparison to the reference Population “A” or “B”, cultured separately, shows adjustment of ‘losing’ population within the “AB” mix. All experimental results (n=9) are shown in Fig. S5.

Fig. 3B shows typical synchronization outcomes (n=32 triplets of RAFL experiments). Strikingly, in almost all experiments (31/32, Fig. S4), the phase of the mixed cell ensemble AB matches very closely the phase of one of the reference cell ensembles, either A or B. This suggests a strongly asymmetric synchronization dynamics: while one oscillator ensemble maintains its rhythm and essentially remains unchanged, the second oscillator ensemble fully adjusts its rhythm to match the “winner’s” phase. We verified through dual-reporter experiments that within the mixed cell ensemble, cells originating from both tailbuds were indeed oscillating and hence verified that the “losing” ensemble acquired the winners’ rhythm (Fig. 3C, Fig. S5). We also verified that both reference cell ensembles were oscillating with a similar period (Fig. S6) to ensure that this synchronization outcome was due to phase differences, not period differences. This “winner-takes-it-all” synchronization result is not compatible with the phase averaging prediction made by the Kuramoto model (Fig. 1D). The discrepancy is especially evident for oscillators initially close to anti-phase, as in that case, the “losing” oscillator ensemble shifts by a phase *π* to lock on the “winning” oscillator ensemble’s phase (Fig. 3B, second row). To further investigate this unexpected winner-takes-it-all synchronization and determine coupling rules compatible with our data, we turned to mathematical modeling.

### Inferring coupling rules for winner-takes-it-all synchronization

The highly asymmetric synchronization that we observe is incompatible with Kuramoto coupling. We thus turned to mathematical modelling to establish a minimal model compatible with data based on phase response theory [19, 25]. For simplicity, we model the behaviour of each oscillator ensemble in the mixed population AB with a single phase variable (*ϕ*_*A*_ for cells coming from embryo A and *ϕ*_*B*_ for cells coming from embryo B). We focus on their coupling, assuming the ensemble oscillators are well-mixed so that we can coarse-grain couplings with functions depending purely on *ϕ*_*A*_, *ϕ*_*B*_. In general, the coupling signal sent by a given oscillator can vary depending on its phase *ϕ* . Similarly, the phase of the oscillator that receives the coupling signal can impact its response to such signal. Such variations of the coupling signal and response can be captured by functions *S*(*ϕ*) and *R*(*ϕ*), respectively [25]. However, if the coupling strength *c* is weak, such that the coupled oscillators’ phases vary slowly compared to the period of oscillations, we can average the system’s behavior over one oscillation cycle, and therefore capture the coupling rules with a single function *H* [26]: with

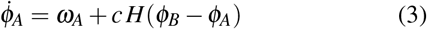

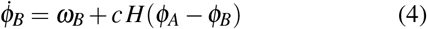

With

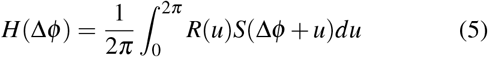

(see the SI Text for a detailed derivation adapted from [19]). By averaging out all specific phase effects into a dependency on the phase difference, a single *H* function can capture many possible forms of the signal function *S* and of the response function *R*.

Next, we constrain the shape of the coupling function *H* so that it is compatible with the winner-takes-it-all dynamics (see our detailed mathematical reasoning in the SI Text). Since one oscillator remains unchanged during synchronization, it means that its coupling term *H*(Δ*ϕ*) has to remain fixed at 0 at all times. For the losing oscillator ensemble though, this coupling term should not be 0 since it adjusts to the other oscillator ensemble. Eventually, the oscillators synchronize such that Δ*ϕ* = 0. This means the *H* function should be asymmetric and exactly 0 for either positive or negative phase differences. However, *H* should also be 2*π* periodic since it is a function of phases. The simplest functions *H* with such properties giving stable in-phase synchronization with a coupling constant *c >* 0 are :

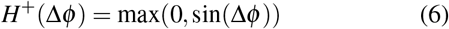

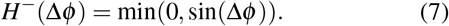

*H*^+^ and *H*^*–*^ are shown in Fig. 4 (A and B). Close to 0, such functions behave like “Rectified Linear Units” or “ReLU” [27], routinely used in machine learning. We thus call such coupling functions “Rectified Kuramoto” or “ReKu”.

**FIG. 4.**
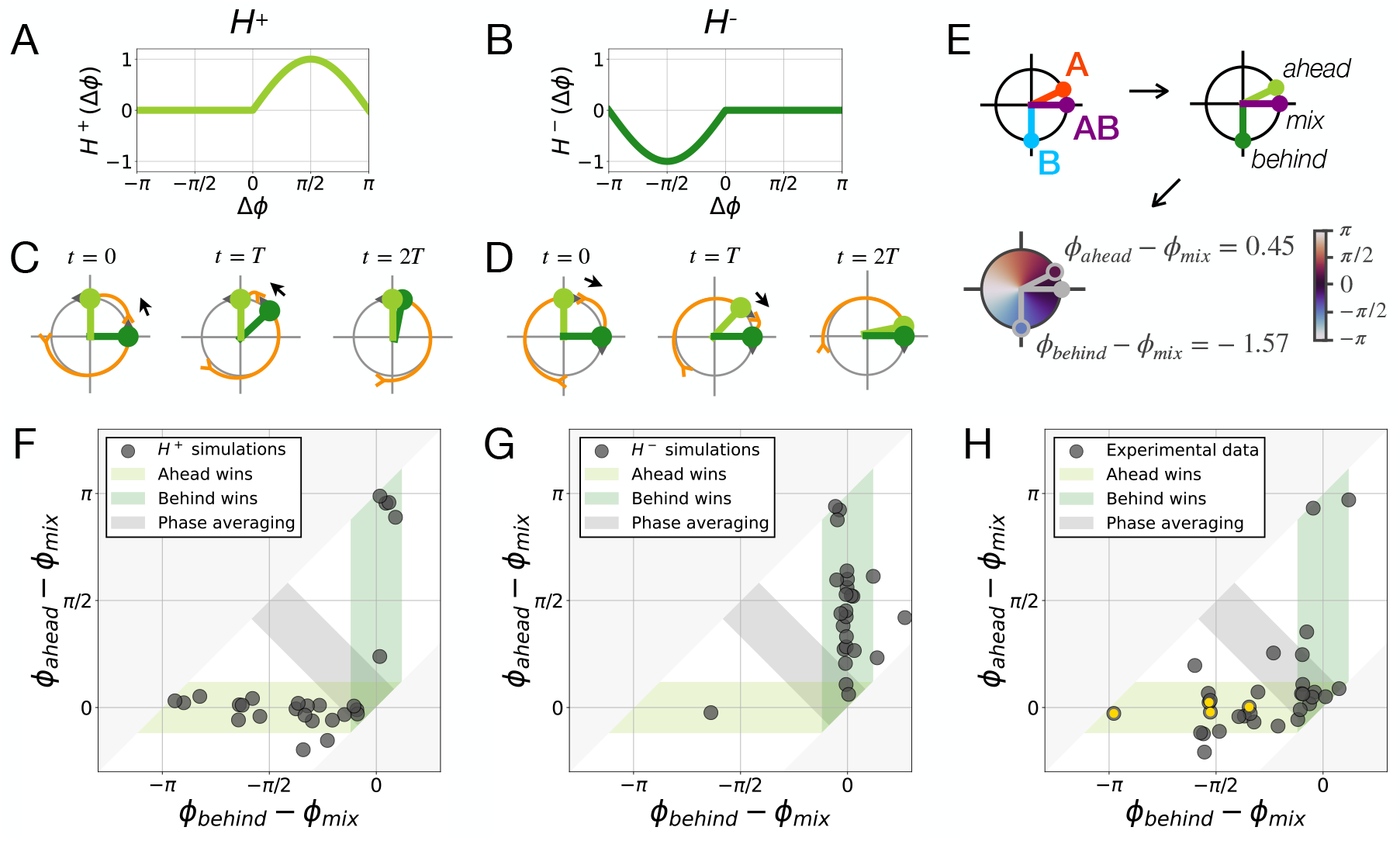
Coupling functions for winner-takes-it-all dynamics. (**A** and **B**) Asymmetric coupling functions *H*^+^ (**A**) and *H*^*–*^ (**B**). (**C** and **D**) Schematic of the synchronization dynamics with coupling functions *H*^+^ (**C**) and *H*^*–*^ (**D**). Either the oscillator ahead (light green) pulls the other oscillator (*H*^+^, **C**) or the oscillator behind (dark green) pulls the other oscillator (*H*^*–*^, **D**). (**E**) Schematic of the data analysis procedure to identify which oscillator is ahead and which is behind, thus defining three phases, *ϕ*_*ahead*_, *ϕ*_*behind*_, and *ϕ*_*mix*_, and their differences after stabilization. (**F**-**H**) Once defined in (**E**), we plot *ϕ*_*ahead*_ – *ϕ*_*mix*_ as a function of *ϕ*_*behind*_ – *ϕ*_*mix*_. In a “winner-takes-it-all” synchronization, one of these phase differences should be 0. Indeed, the horizontal axis, where *ϕ*_*ahead*_ – *ϕ*_*mix*_ = 0, corresponds to a situation where the “ahead” oscillator wins, and the vertical axis, where *ϕ*_*behind*_ –*ϕ*_*mix*_ = 0, corresponds to a situation where the “behind” oscillator wins, while the diagonal corresponds to phase averaging. To set the width of each colored region, we estimated the phase difference between synchronized oscillators expected from experimental limitations by calculating the standard deviation of the ROI phases for each RAFL experiment with more than one ROI (n=82). (**F**) Numerical simulations of the two-way synchronization experiment Eqs. 3-4 with *H* = *H*^+^. We performed 32 simulations with random initial phases and random periods that model the data’s phase and period distributions. Points concentrate on the horizontal axis showing that “ahead” oscillators are winning. (**G**) Numerical simulations of the two-way synchronization experiment Eqs. 3-4 with *H* = *H*^*–*^. Points concentrate on the vertical axis showing that “behind” oscillators are winning. (**H**) Experimental results of the two-way synchronization experiments. Phases are measured according to the procedure defined in panel (**E**), at *t* = 400 min. Yellow dots highlight the experiments shown in Fig. 3B. Points are concentrated on the horizontal axis, suggestive of a *H*^+^ coupling.

### Double asymmetry for ReKu coupling

The ReKu coupling functions achieve winner-takes-it-all synchronization because they possess a double asymmetry: a response only occurs in half of the cycle, and in addition, its direction is of constant sign i.e., either always positive (Fig. 4A) or always negative (Fig. 4B). The first asymmetry ensures that one oscillator remains unchanged despite being coupled to the other oscillator. The second asymmetry makes the loser adjust its rhythm to match the winner’s phase, either by speeding up (Fig. 4C, arrow) or by slowing down (Fig. 4D, arrow). Furthermore, because the ReKu coupling functions can only reduce the phase difference between the two oscillators, the second asymmetry also entails that either the oscillator initially ahead is always winning (Fig. 4C and F), or the oscillator initially delayed is always winning (Fig. 4D and G). To evaluate this prediction, we returned to the experimental data.

### Ahead oscillator is winning, suggestive of *H*^+^ coupling

To distinguish whether *H*^+^ or *H*^*–*^ coupling rules predict the synchronization outcome in the experiments, we analyzed how the initial phase relation between the ensembles would correlate with the synchronization outcome (Fig. 4E-H). We found that the oscillator initially ahead was winning most of the time (27 out of 32 triplets of RAFL experiments, Fig. 4H). Importantly, being “ahead” was predictive of winning for a wide range of initial phases, hence suggesting that being ahead, rather than being close to any specific phase, is essential. For completeness, we also investigated the 5 out of 32 apparent outliers. They can be accounted for by either a slight mismatch in the intrinsic frequencies (Fig. S7A), or by ambiguity in labeling which oscillator is ahead when oscillators are initially close to anti-phase (Fig. S7B). Our results hence suggest that randomized tailbud cells synchronize with a ReKu coupling function of the *H*^+^ form.

### Evaluating alternative coupling models

While the ReKu coupling model explains our synchronization data and predicts that the ahead oscillator ensemble should always win, it is also important to confront data with alternative coupling models, to see if they can equally account for the winner-takes-it-all outcome. First, we examined the Kuramoto-Sakaguchi (KS) coupling function [28]:

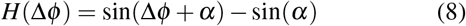

This coupling term translates the Kuramoto coupling function by a phase *α*, while ensuring that *H*(0) = 0 so that oscillators initially in phase stay in phase (see arrows in Fig. 5A). In contrast to the Kuramoto coupling function, the KS coupling introduces an asymmetry in *H*, for instance, for *α* = *–*1, the KS coupling function is positive almost everywhere, and thus mostly speeds up oscillators. However, the KS coupling function is lacking the second asymmetry present in the ReKu coupling, i.e. the response is not restricted to half the cycle. As a result, while for some specific *α ∼ ±*1, winner-takes-it-all synchronization can be obtained for most phase shifts Δ*ϕ* (Fig.5B), for phase shifts closer to *π*, the synchronization outcome strongly diverges from winner-takes-it-all (Fig. S8A), which is incompatible with our data (Fig. 3, second row; Fig. S4). In the SI Text, we further show that delayed coupling models [21, 29] can be reduced to KS coupling, with the *α* parameter depending on the delay *τ*, the intrinsic frequency Ω and the coupling strength *c*. Therefore, we conclude that the KS model, and thus delayed coupling, cannot explain the experimentally observed winner-takes-it-all synchronization.

**FIG. 5.**
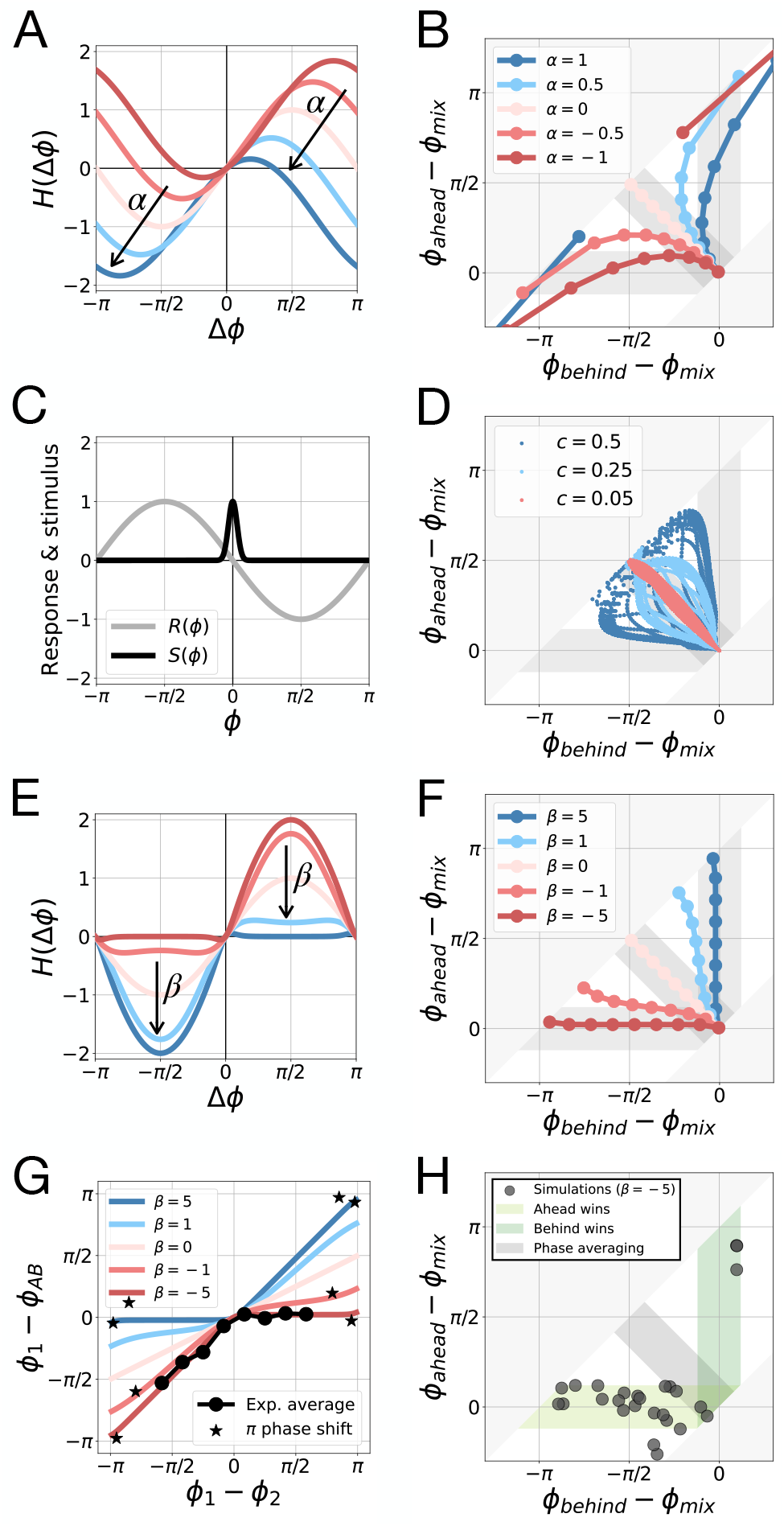
Testing alternative models. (**A**) Coupling function for the Kuramoto-Sakaguchi model with different values of parameter *α*. (**B**) Simulation results of two-way synchronization for the Kuramoto-Sakaguchi model. (**C**) Response and stimulus functions for the pulsed-coupling model. (**D**) Simulation results of two-way synchronization for the pulsed-coupling model with different coupling strengths *c*. (**E**) Coupling function for the continuous ReKu model with different values of parameter *β* . (**F** and **G**) Simulation results of two-way synchronization for the continuous ReKu model, compared to experimental data (**G**). *ϕ*_1_ represents either *ϕ*_*A*_ or *ϕ*_*B*_. If *ϕ*_1_ = *ϕ*_*A*_, then *ϕ*_2_ = *ϕ*_*B*_, and vice versa. In experiments with a *π* phase shift between *ϕ*_*A*_ and *ϕ*_*B*_, determining unambiguously which oscillator is ahead is impossible. For this reason, these experiments were not included in the data average. (**H**) Simulation results for the ReKu model with *β* = *–*5. We performed 32 simulations with random initial phases and random periods that model the data’s phase and period distributions.s

Next, we investigated “pulsed-coupling” models. For such coupling models, an oscillator emits a strong signal only when reaching a given phase (signal function *S*(*ϕ*) on Fig. 5C). This leads to an intermittent (rather than continuous) synchronization response from the second oscillator (response function *R*(*ϕ*) on Fig. 5C). As explained above, such signal and response functions are not captured by a single H function if the coupling strength *c* is strong (for weak coupling, pulsed-coupled models can be described by an effective H function). We thus tested if pulsed-coupling models in the strong coupling regime could achieve winner-takes-it-all synchronization (Fig. 5D). We found that winner-takes-it-all dynamics were possible, but only for very strong coupling (*c ∼* 10 *ω*) and importantly, only for small phase shifts (Fig. 5D, Fig. S8C). Accordingly, for initial phase shifts close to *π* the synchronization outcome strongly diverges from winner-takes-it-all, contrary to experiments (Fig. S8C). We conclude that pulsed-coupling models in the strong coupling regime cannot account well for the synchronization outcomes we observed. We also investigated in more detail the impact of asymmetries (Fig. 5E) and additional coupling models with different asymmetries (Fig. S9) on the synchronization outcome. To examine the transition from models without asymmetry, such as the standard Kuramoto model, to ReKu models with a double asymmetry, we considered the following *H* function:

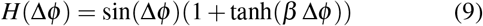

which continuously changes with parameter *β* from the ReKu model *H*^+^ (*β → –*∞), to the Kuramoto model (*β* = 0), and then to the ReKu model *H*^*–*^ (*β →* +∞) (Fig. 5E). We found that only coupling functions with an almost complete double asymmetry (*β << –*1) result in a winner-takes-it-all synchronization compatible with data (Fig. 5, F and G). We further generated synthetic data by simulating a ReKu model with *β* = *–*5, random initial phases, and a small noise on the intrinsic frequencies (Fig. 5H). The resulting synthetic data is very similar to the experimental data: in most simulations, the oscillator initially ahead is winning. In the remaining few simulations, the oscillator initially behind appears to be winning, because of a specific combination of initial phases and frequency mismatch (Fig. S7). We conclude that winner-takesit-all synchronization requires the coupling function to possess a double asymmetry similar to the ReKu models.

## DISCUSSION

In this work, our goal was to identify the coupling rules within an embryonic oscillator ensemble. We combined theoretical modeling with a novel experimental strategy that allows direct determination of the emergent phase of synchronization. Critically, using single embryos as input in the RAFL assays, we were able to compare the collective, emergent phase reached after synchronization in the intermixed cell ensemble with the reference phase of each individual input cell ensemble, for a broad distribution of initial phases. We found that one oscillator remains essentially unaltered in its phase dynamics, while the other population is adjusting its phase to synchronize with the former. Such an asymmetric “winner-takes-it-all” synchronization outcome is incompatible with the Kuramoto model. We therefore devised a new model, the Rectified Kuramoto (ReKu) model. The ReKu model achieves winner-takes-it-all synchronization thanks to its double asymmetry: the response is always positive and restricted to only parts of the cycle. As an effect, the oscillator “ahead” is always winning.

Linking our results to the dynamics seen in the natural *in vivo* context, it is well established that PSM cells exhibit a slowing down of their oscillations over time [13, 30]. The consequence is a posterior-to-anterior gradient of increasing period that is a hallmark of this dynamical system and underlies the emergence of phase waves that sweep across the embryo axis. At first, the slowing down seen *in vivo* appears to be at odds with our observation that tailbud cells used in the RAFL assay can only speed up their oscillations when coupled. One interpretation that might follow is that coupling and the input signal is modulated along the antero-posterior embryo axis, thereby ensuring both synchronization in the tailbud cells (via speeding up and winner-takes-it-all dynamics) and subsequent slowing down in the PSM. A recent study using cells from the entire PSM (in contrast to the tail bud cells we used in RAFLs) did find evidence for a constant sign response, yet the effect of coupling was a delay of oscillations [31]. The more general insight from studying PSM oscillators *in vitro* is hence that of a highly asymmetric coupling mechanism that results in a constant sign response. Interestingly, recent entrainment experiments of segmentation clock oscillators performed on embryonic PSM tissue explants revealed a similar, highly asymmetric response, in this case to periodic entrainment pulses [32]. In these experiments, the segmentation clock oscillator’s phase response curve (PRC) to pulses of a Notch inhibitor (DAPT) was found to be close to 0 for a significant part of the cycle, and mostly negative otherwise, meaning that DAPT mostly delays the segmentation clock. The qualitatively similar asymmetric responses in both entrainment and RAFL experiments might reflect the intrinsic geometry and bifurcation structure of the segmentation oscillator itself.

Accordingly, phase responses of constant sign generally appear in systems poised near an infinite-period bifurcation [33] such as the saddle-node on invariant cycle (SNIC) bifurcation [34]. The normal form of oscillators close to SNICs correspond to the “Integrate and fire” model [33]. Such models were used to explain synchronization and wave propagation in other contexts e.g., for the emergence of light rhythms and synchronization of fireflies [35, 36]. Importantly, one does not need the full knowledge of internal mechanisms (e.g. gene interactions) to describe, understand, and predict such oscillatory patterns. Fundamentally, this comes from the fact that (the change of) behaviours can be systematically classified using tools from bifurcation and catastrophe theory [37, 38]. Recent studies have further shown how gene-regulatory networks based on cell differentiation data can be conceptualized into “landscapes”, leading to the description of global bifurcations relevant for decision-making in biology such as the “heteroclinic flip” [39], allowing for a richer set of behavior and control to drive cellular fates [40]. The asymmetric shape of oscillator coupling observed here is suggestive of the proximity to infinite period bifurcations, which are global bifurcations for oscillators (by opposition to local Hopf bifurcations that would result in Kuramoto coupling), and thus appear to drive phase decision of oscillators. Indeed such asymmetries of couplings are already known to be used in complex systems to perform sophisticated computations, from neural couplings [33] to size-effects of human clapping synchronization [41]. An infinite period bifurcation allows for efficient modulations of period and phase with changes of control parameters [33], providing a possible explanation for its function in an embryonic oscillator ensemble.

## MATERIALS AND METHODS

More detailed experimental methods and mathematical derivations are available in the Supplementary Materials. All codes used for this work are available at the following url: https://github.com/laurentjutrasdube/Nonreciprocal_synchronization_in_embryonic_oscillator_ensembles/

## ACKNOWLEDGEMENTS

We thank all members of the A.A. and P.F. groups for their important input and insights. This work was supported by the European Molecular Biology Laboratory and received funding from the European Research Council under an ERC consolidator grant agreement n.866537 to A.A. This work is supported by the EMBL Advanced Light Microscopy Facility (ALMF) and EMBL Laboratory Animal Resource (LAR). This work was supported by a Bridging Excellence Fellowship provided by the Life “Science Alliance to M.Z.This work was supported by Simons Investigator in Mathematical Modeling of Living Systems, NSERC Discovery grant, CIHR Program Grant, and Fonds Courtois awards to P.F.; a Fonds de recherche du Québec–Nature et Technologies doctoral scholarship and a National Science and Engineering Research Council-CREATE in Complex Dynamics graduate scholarship to L.J.D.

