## Supplementary Information for "Nonreciprocal synchronization in embryonic oscillator ensembles"

#### This PDF file includes:

Supporting text
Figs. S1 to S9
Table S1
Legends for Movies S1 to S2
SI References

#### Other supporting materials for this manuscript include the following:

Movies S1 to S2

### Supporting Information Text

**From the generic Winfree model of phase oscillator coupling to Kuramoto's phase-difference dependent coupling function.** We start with the following differential equations to describe the phase dynamics of coupled oscillators $A$ and $B$ (1):

$$\begin{aligned}\dot{\phi}_A &= \omega_A + c\, Z(\phi_A)\, \Gamma(\phi_A, \phi_B)\\ \dot{\phi}_B &= \omega_B + c\, Z(\phi_B)\, \Gamma(\phi_B, \phi_A)\end{aligned} \quad [1]$$

where $c$ represents the coupling strength, and $Z$ describes the response of each oscillator to the external perturbation $\Gamma$ imposed by the other oscillator. We assume that the coupled oscillators reach a common frequency $\Omega$, and perform the following change of variables:

$$\begin{aligned}\phi_A &= \Omega\, t + \Psi_A\\ \phi_B &= \Omega\, t + \Psi_B\end{aligned} \quad [2]$$

Injecting this Ansatz into Eq. 1 gives the differential equations for $\Psi_A$ and $\Psi_B$:

$$\begin{aligned}\dot{\Psi}_A &= \omega_A - \Omega + c\, Z(\Psi_A + \Omega\, t)\, \Gamma(\Psi_A + \Omega\, t, \Psi_B + \Omega\, t)\\ \dot{\Psi}_B &= \omega_B - \Omega + c\, Z(\Psi_B + \Omega\, t)\, \Gamma(\Psi_B + \Omega\, t, \Psi_A + \Omega\, t)\end{aligned} \quad [3]$$

This change of variable allows us to simplify the coupling terms. Indeed, if $\Psi_A$ and $\Psi_B$ vary slowly, we can safely average Eq. 3 over one period $T = 2\pi/\Omega$ to get the following equations:

$$\begin{aligned}\dot{\Psi}_A &= \omega_A - \Omega + c\, H(\Psi_B - \Psi_A)\\ \dot{\Psi}_B &= \omega_B - \Omega + c\, H(\Psi_A - \Psi_B)\end{aligned} \quad [4]$$

where

$$H(\Psi_B - \Psi_A) = \frac{1}{T}\int_0^T Z(\Psi_A + \Omega t)\, \Gamma(\Psi_A + \Omega t, \Psi_B + \Omega t)\, dt = \frac{1}{2\pi}\int_0^{2\pi} Z(u)\, \Gamma(u, \Psi_B - \Psi_A + u)\, du \quad [5]$$

where we performed a change of variable $u = \Omega t + \Psi_A$ to eliminate the $\Omega$ dependency, using the fact that $Z$ and $\Gamma$ are $2\pi$-periodic in their phase arguments to perform the period averaging. This procedure renders explicit the dependency on the phase difference $\Psi_B - \Psi_A$. Since $\Psi_B - \Psi_A = \phi_B - \phi_A$, we can also go back to the initial variables:

$$\begin{aligned}\dot{\phi}_A &= \omega_A + c\, H(\phi_B - \phi_A)\\ \dot{\phi}_B &= \omega_B + c\, H(\phi_A - \phi_B)\end{aligned} \quad [6]$$

which are Eq. 3 & 4 given in the main text. It is of course always possible to study directly such system of equations, but notice it is only equivalent to the full version when both oscillators are assumed to lock to a common frequency. In his 1984 paper, Kuramoto studied the synchronization dynamics of Eq. 6 with $H(\Delta\phi) = \sin(\Delta\phi)$ (2). A follow-up paper published 2 years later with Sakaguchi extended the analysis to sine functions with any offset $\alpha$: $H(\Delta\phi) = \sin(\Delta\phi + \alpha)$ (3).

**Construction of the rectified Kuramoto (ReKu) coupling function.** To determine which properties the $H$ function must have to produce winner-takes-it-all synchronization, we first note that the $H$ function directly governs the temporal evolution of a given coupled oscillator relative to its reference oscillator. Indeed, since the temporal evolution of the reference oscillators is respectively given by $\dot{\phi}^r_A = \omega_A$ and $\dot{\phi}^r_B = \omega_B$, we have:

$$\frac{d}{dt}(\phi_A - \phi^r_A) = c\, H(\phi_B - \phi_A) \quad [7]$$

$$\frac{d}{dt}(\phi_B - \phi^r_B) = c\, H(\phi_A - \phi_B) \quad [8]$$

The winner-takes-it-all dynamics means that at least one of those equations is always 0, irrespective of the (changing) values of $\phi_A$ and $\phi_B$. Formally, setting $\Delta\phi = \phi_A - \phi_B$, this means that

$$H(\Delta\phi)\, H(-\Delta\phi) = 0 \quad [9]$$

for all values of $\Delta\phi$ taken experimentally. Eq. 9 implies that $H(\Delta\phi = 0) = 0$, which is consistent with the fact that oscillators eventually synchronize with no change of intrinsic frequencies (Fig. S4A). Furthermore, Eq. 9 suggests that $H$ has a singular, asymmetric behaviour close to 0. To see this, let us assume that $H$ is differentiable in 0, meaning that sufficiently close to $|\Delta\phi| = 0$, $H(\Delta\phi) \sim A\Delta\phi^n$ where $n$ is the order of the first non-zero derivative close to 0. Then, Eq. 9 for non-zero $\Delta\phi$ imposes that $A$ is 0, which implies that $H = 0$ everywhere. To solve this conundrum, the simplest solution is to assume that $H$ is not differentiable in 0, and with Eq. 9 the simplest hypothesis is to assume that $H$ is linear on one side and identically 0 on the other side. There are thus four possible qualitative behaviours depending on the sign of the derivative in 0 and the choice of the non-zero side of $H$. Only two of these four choices give stable synchronized coupling:

$$H(\Delta\phi) = \begin{cases} 0 & \text{for} \quad \Delta\phi < 0 \\ a\Delta\phi & \text{for} \quad \Delta\phi > 0 \end{cases} \quad \text{and} \quad H(\Delta\phi) = \begin{cases} a\Delta\phi & \text{for} \quad \Delta\phi < 0 \\ 0 & \text{for} \quad \Delta\phi > 0 \end{cases} \quad [10]$$

where $a > 0$ ensures that $\Delta\phi = 0$ is stable, which means that synchronized oscillators stay synchronized. Periodic generalization of this rectified linear behaviour close to 0 is achieved by considering only the rectified part of the Kuramoto coupling function i.e., taking:

$$H^{+}(\Delta\phi) = a\max(0,\, \sin(\Delta\phi)) \qquad \text{or} \qquad H^{-}(\Delta\phi) = a\min(0,\, \sin(\Delta\phi)) \tag{11}$$

as indicated in Eq. 6 and 7 of the main text.

**Delayed coupling and the Kuramoto-Sakaguchi model.** The goal of this section is to show that, in our theoretical framework, models based on delayed Kuramoto coupling are equivalent to the Kuramoto-Sakaguchi model. Following (4), we consider phase oscillators coupled via the Kuramoto coupling function with a delay:

$$\dot{\phi}_i = \omega_i + c\sum_j \sin\big(\phi_j(t-\tau) - \phi_i(t)\big) \tag{12}$$

where $\tau$ accounts for various biochemical delays. Considering two coupled oscillators with a common frequency $\omega$, we get:

$$\begin{aligned} \dot{\phi}_A &= \omega + c\sin\big(\phi_B(t-\tau) - \phi_A(t)\big) \\ \dot{\phi}_B &= \omega + c\sin\big(\phi_A(t-\tau) - \phi_B(t)\big) \end{aligned} \tag{13}$$

To study the coupling dynamics, we assume that both oscillators end up oscillating with a common frequency $\Omega$, and we define:

$$\begin{aligned} \phi_A(t) &= \Omega\, t + \Psi_A \\ \phi_B(t) &= \Omega\, t + \Psi_B \end{aligned} \tag{14}$$

where $\Psi_A$ and $\Psi_B$ respectively define the relative phases of oscillators $A$ and $B$ close to stationarity (i.e. they vary very slowly compared to $\Omega$, similar to what is done in the previous section). The differential equations for $\Psi_A$ and $\Psi_B$ are:

$$\begin{aligned} \dot{\Psi}_A &= \omega - \Omega + c\sin(\Psi_B - \Psi_A - \Omega\tau) \\ \dot{\Psi}_B &= \omega - \Omega + c\sin(\Psi_A - \Psi_B - \Omega\tau) \end{aligned} \tag{15}$$

Taking the difference, at stationarity we get:

$$\begin{aligned} 0 &= \sin(\Psi_B - \Psi_A - \Omega\tau) - \sin(\Psi_A - \Psi_B - \Omega\tau) \\ \implies\ 0 &= \sin(\Psi_B - \Psi_A)\cos(\Omega\tau) \end{aligned} \tag{16}$$

In general, Eq. 16 implies that for positive $c$, the only stable phase relation is $\Psi_B - \Psi_A = 0 \mod 2\pi$. Injecting this phase relation into equation 15, we get (4):

$$\Omega = \omega - c\sin(\Omega\tau) \tag{17}$$

This is a transcendental equation and there is no analytical solution, but, assuming that $\tau$ is small, we get the following approximate expression:

$$\Omega \sim \frac{\omega}{1 + c\tau} \tag{18}$$

This shows that if the delay is high enough, the coupled oscillators go slower than their "intrinsic" frequency $\omega$.

Now, we inject Eq. 17 into Eq. 13:

$$\begin{aligned} \dot{\phi}_A &= \Omega + c\Big[\sin\big(\phi_B(t-\tau) - \phi_A(t)\big) + \sin(\Omega\tau)\Big] \\ \dot{\phi}_B &= \Omega + c\Big[\sin\big(\phi_A(t-\tau) - \phi_B(t)\big) + \sin(\Omega\tau)\Big] \end{aligned} \tag{19}$$

Finally, we use Eq. 14 close to stationarity, i.e. assuming that $\Psi_A$ and $\Psi_B$ are almost constant ($\Psi_A(t) \sim \Psi_A(t-\tau) \sim \Psi_A$), to write :

$$\begin{aligned} \phi_A(t-\tau) &= \Omega(t-\tau) + \Psi_A = \phi_A(t) - \Omega\tau \\ \phi_B(t-\tau) &= \Omega(t-\tau) + \Psi_B = \phi_B(t) - \Omega\tau \end{aligned} \tag{20}$$

which allows us to get our final result:

$$\begin{aligned} \dot{\phi}_A &= \Omega + c\Big[\sin\big(\phi_B(t) - \phi_A(t) - \Omega\tau\big) - \sin(-\Omega\tau)\Big] \\ \dot{\phi}_B &= \Omega + c\Big[\sin\big(\phi_A(t) - \phi_B(t) - \Omega\tau\big) - \sin(-\Omega\tau)\Big] \end{aligned} \tag{21}$$

Therefore, close to stationarity, introducing a delay $\tau$ in the Kuramoto coupling function corresponds to using the Kuramoto-Sakaguchi coupling function, $H(\Delta\phi) = \sin(\Delta\phi + \alpha) - \sin(\alpha)$, with a value of $\alpha = -\Omega\tau$ (see Eq. 8 of the main text). Using the parameters from (4), i.e a delay of $\tau = 7$ min, and a collective period of $T = 39.1$ min, corresponding to a collective frequency of $\Omega = 2\pi/T = 0.161\ \text{min}^{-1}$, we get a value of $\alpha = -1.13$. As explained in the main text, the Kuramoto-Sakaguchi model with values of $\alpha$ close to $-1$ display winner-takes-it-all dynamics for a large interval of initial conditions for the two coupled oscillators' phases (Fig. 5B), but fails to generate winner-takes-it-all dynamics for initial phase differences close to $\pi$ (Fig. S8A), in contrast with the RAFL data (Fig. 3B, second row, and Fig. S5B).

**Pulsed-coupling models and piece-wise linear versions.** To build the pulsed-coupling model, we use Winfree's theoretical framework (see (1), and the first section of this SI Text) with two coupled oscillators $\phi_A$ and $\phi_B$:

$$\dot{\phi}_A = \omega + c\, S(\phi_B)\, R(\phi_A) \qquad [22]$$

$$\dot{\phi}_B = \omega + c\, S(\phi_A)\, R(\phi_B) \qquad [23]$$

with a Gaussian signal function $S$:

$$S(\phi) = e^{-\phi^2 / 2\sigma^2} \qquad [24]$$

and a sine response function $R$:

$$R(\phi) = \sin(-\phi) \qquad [25]$$

In the rectified pulsed-coupling model, we replace the response function $R$ by an expression similar to Eq. 9 of the main text:

$$R(\phi) = \sin(-\phi)\Big(1 - \tanh\big(\beta \sin(-\phi)\big)\Big) \qquad [26]$$

In the piece-wise linear rectified pulsed-coupling model, we replace the response function $R$ by the following function:

$$R(\phi) = \begin{cases} -\phi\,\big(1 - \gamma\,\mathrm{sign}(\phi)\big) & \text{for} |\phi| < \frac{\pi}{2} \\ (\pi\,\mathrm{sign}(\phi) + \phi)\,\big(1 - \gamma\,\mathrm{sign}(\phi)\big) & \text{for} |\phi| > \frac{\pi}{2} \end{cases} \qquad [27]$$

where parameter $\gamma$ plays the role of parameter $\beta$ (Eq. 26). Similarly, in the piece-wise linear rectified Kuramoto model, the $H$ function of ODEs:

$$\dot{\phi}_A = \omega + c\, H(\phi_B - \phi_A) \qquad [28]$$

$$\dot{\phi}_B = \omega + c\, H(\phi_A - \phi_B) \qquad [29]$$

becomes:

$$H(\Delta\phi) = \begin{cases} \Delta\phi\,\big(1 - \gamma\,\mathrm{sign}(\Delta\phi)\big) & \text{for} |\Delta\phi| < \frac{\pi}{2} \\ (\pi\,\mathrm{sign}(\Delta\phi) - \Delta\phi)\,\big(1 - \gamma\,\mathrm{sign}(\Delta\phi)\big) & \text{for} |\Delta\phi| > \frac{\pi}{2} \end{cases} \qquad [30]$$

where again, parameter $\gamma$ plays the role of parameter $\beta$ (Eq. 9 of the main text).

### Methods

**Mouse lines.** The majority of the experiments were performed using a transgenic mouse line expressing a dynamic Notch signaling reporter controlled by the Lfng promoter and also have a constitutive expression of H2B-mCherry. This mouse line is named: H2BmCherry/LuVelu. The generation of the LuVelu mouse line was previously described in (5).

**Medium preparation.** Dissection and culture media were freshly prepared as indicated in Table S1: culture medium was filtered using a PVDF filter, pore size 0.22 µm (Merck) and kept in the incubator at 37 °C for at least 15 minutes to equilibrate it.

**Randomization assay.** To perform randomization experiments, we used the H2BmCherry/LuVelu mouse line. Mouse embryos were collected at 10.5 dpc (days post coitum) in a dissection medium containing DMEM/F-12 (Cell Culture Technologies) supplemented with 2 µM of Glucose, 2 µM L-Glutamine (Thermofisher 25030081), 0,5 % of BSA, HEPES and penicillin/Streptomycin (Thermofisher 15140122). Using a scalpel, PSM tissues were cut at the very posterior tip of the mouse tails to ensure a narrow phase and frequency distribution between cells. Randomization process included mechanical dissociation, cell filtration through a 30 µm filter (ParTec) and reaggregation of the PSM tissue in the micro insert 4 well Ful trac (Ibidi 80486) coated with fibronectin (Sigma-Aldrich F1141). Finally, PSM randomized cells were cultured overnight at 37 °C and 5 % CO2, in DMEM/F-12 medium supplemented with 2 µM of Glucose, 2 µM L-Glutamine and 0.5 % of BSA.

**Phase synchronization assay.** To study the phase synchronization dynamics, we used two very posterior PSM tissues from two different mouse embryos. Each of them was randomized as described above. One part of each population was kept as a population reference while the other part was mixed in the ratio of 1:1 with the other cell population. To distinguish the origin of two different cell populations, one population carried mCherry as reporter associated with H2B, while the other one didn't.

**Video-microscopy.** Imaging was performed using a Zeiss LSM780 laser-scanning microscope featuring an incubator chamber (EMBL Mechanical Workshop) for CO2 and temperature control. Samples were excited with an Argon laser at 514 nm for Venus and for mCherry. An 20X apo objective was used. Every 5 or 10 minutes a stack of 4 planes at 6 µm was scanned. Multiple samples were recorded using a motorized stage during each experiment. Movies were recorded in 512 × 512 pixels, 1.38 µm per pixel.

**Time series and phase extraction.** After each experiment, movies were collected and analysed with the Fiji software (6). First, the max intensity was extracted using Fiji's functions: Image/Stacks/Z projection and "Max intensity". Then, to make the imaging data smoother, we applied a Gaussian blur filter using a Sigma Radius equal to 10. Time series were then extracted within specific regions of interest (ROI). These ROIs correspond to a miniature emergent PSM (5) formed within a re-aggregate of randomized PSM cells. For each assay, oscillations at the global level were also measured and compared to the calculated average from all smaller ROIs. The global signal and the average ROI signal gave similar time series, with the average signal having a greater oscillation amplitude around the baseline. For this reason, we selected the average data for further analysis of the time series. For each experiment, the average data was detrended using the pyBoat software (7). The same software was used to extract the phase of the average data via wavelet analysis (7).

**Kuramoto order parameter.** For each RAFL experiment with more than one ROI, we computed the Kuramoto order parameter (KOP) as follows:

$$\mathrm{KOP}(t) = \left| \frac{1}{N} \sum_{k} e^{i\,\phi_k(t)} \right| \qquad [31]$$

where $k$ indexes the RAFL experiment's ROIs, from 1 to $N$.

**Code.** All the codes used for this article are available at the following URL :https://github.com/laurentjutrasdube/Nonreciprocal_synchronization_in_embryonic_oscillator_ensembles/

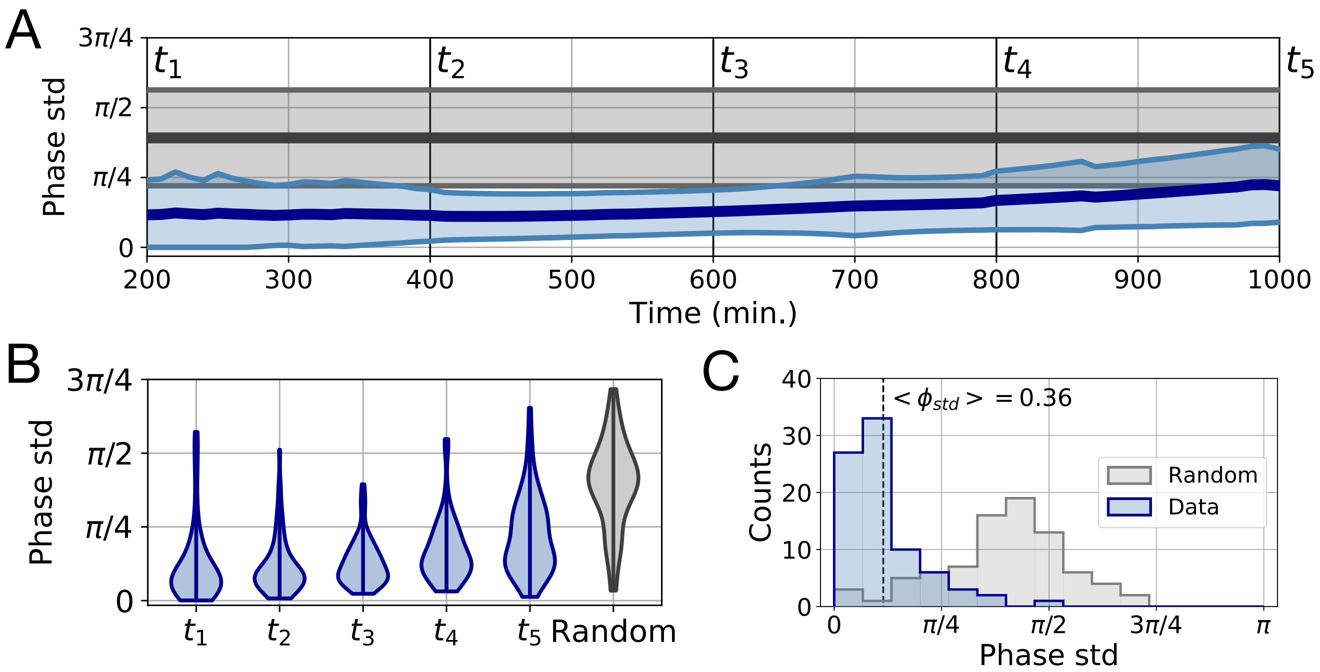


**Fig. S1.** Standard deviation of ROI phases across the RAFL experiments. (**A**) Statistics of the phase standard deviation (std), similarly to the statistics of the Kuramoto order parameter shown in Fig. 2H. One std is computed for every RAFL experiment with more than one ROI (n=82). The thick dark blue line shows the mean std and the thin light blue lines show $\pm$ std. The grey lines show the statistics of the std of random phases with the same distribution as the experiments, i.e. the total number of std computed is equal to the number of experiments, and the number of random phases picked to compute each std is the same as the number of ROIs for the corresponding experiment. (**B**) Violin plots showing the experimental std distribution at each time point identified in (**A**) and the KOP distribution of random phases. (**C**) Phase std distribution of the data at $t_2 = 400$ min. (blue) and of random phases (grey). The data's mean phase std, which provides an estimation of the experimental phase variation, is indicated with a dashed line.

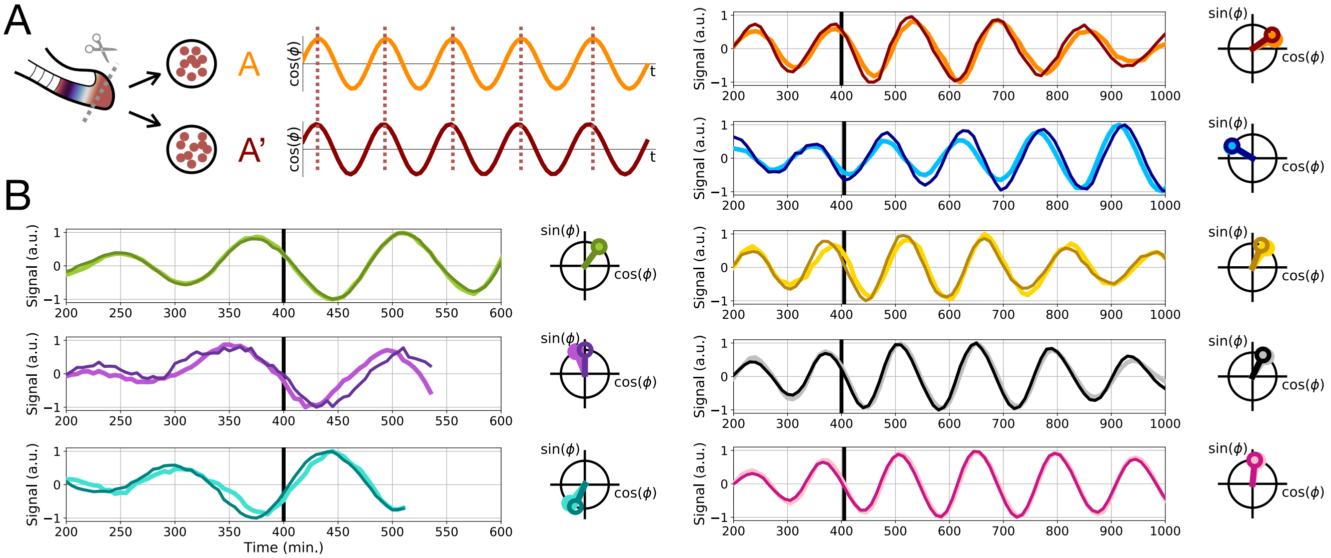


**Fig. S2.** Twin experiments. (**A**) Schematic of the twin experiments: two RAFLs are independently performed with cells coming from the same embryo. (**B**) Time series and polar plots for all twin experiments (n=8). The thick black line indicates the time of the associated polar plot.

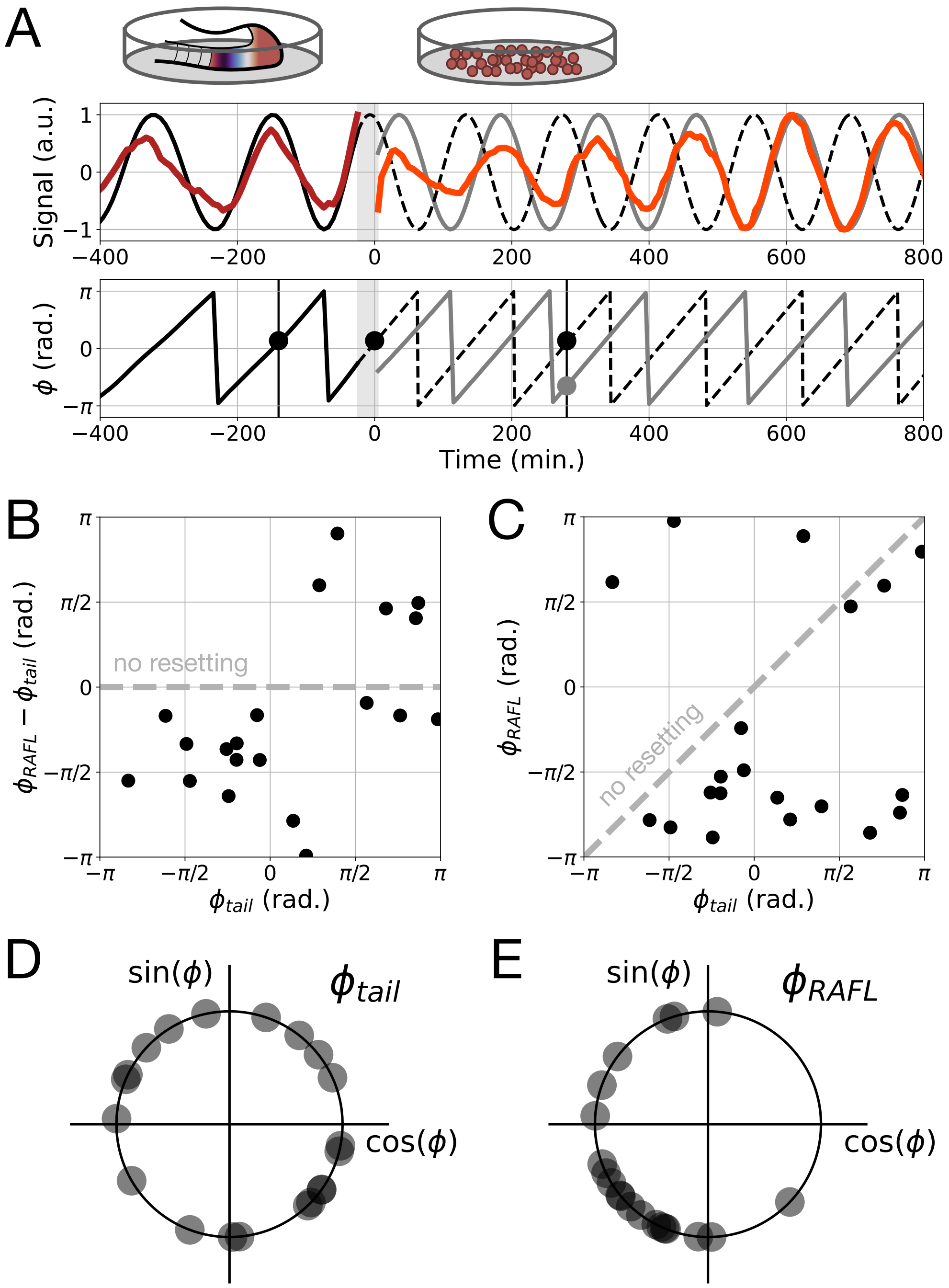


**Fig. S3.** Tail experiments and phase resetting. (**A**) The full PSM is placed in a dish and its oscillations are monitored (dark red), before a RAFL experiment is performed with cells from the PSM's tailbud (orange). The tail and RAFL oscillation phases, resp. in solid black and grey, are extracted using the pyBOAT software. To estimate the resetting due to the RAFL procedure, we compare the phase at the moment when the cells are placed in the incubator (black dot, $\phi_{tail}$) to the RAFL phase two periods later (grey dot, $\phi_{RAFL}$). Since we cannot image the cells as they are placed in the incubator, we estimate $\phi_{tail}$ by computing the tail phase one period before incubation. We set the period to $T = 140$ min. (**B-C**) Phase response (**B**) and transition (**C**) curves (n=20). The dashed grey lines indicate the results expected in the absence of resetting. (**D-E**) Distributions of $\phi_{tail}$ (**D**) and $\phi_{RAFL}$ (**E**) for all tail experiments (n=20). The $\phi_{tail}$ distribution is uniform, while the $\phi_{RAFL}$ distribution is not, consistent with the results shown in Fig. 2J.

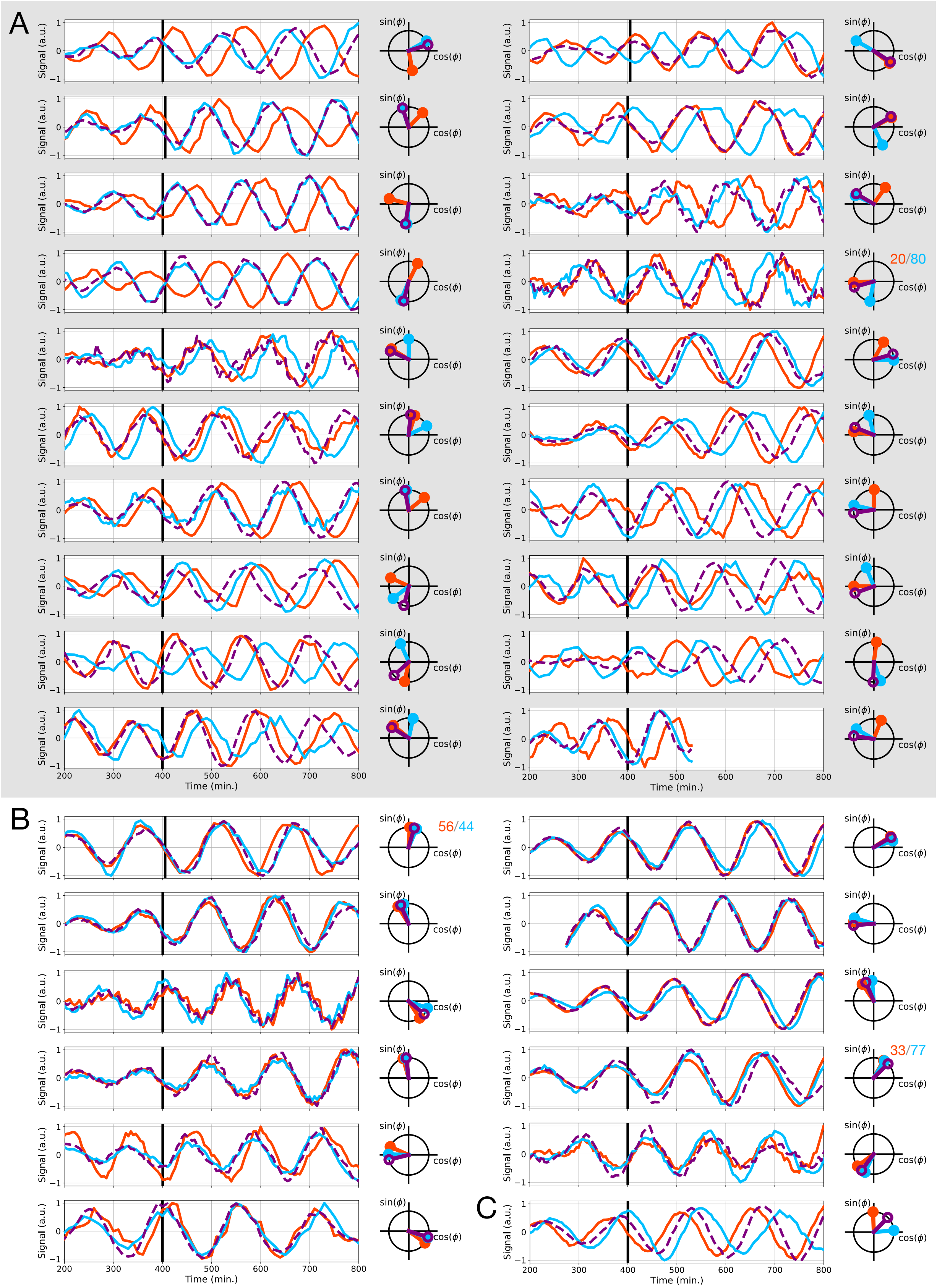


**Fig. S4.** All synchronization experiments (n=32). (**A**) Experiments with winner-takes-it-all dynamics (n=20). The reference population from embryo A is shown in orange, the reference population from embryo B is shown in blue, and the mixed population AB is shown in purple. The thick black line indicates the time of the associated polar plot. The proportion of cells from embryo A (resp. B) in the mix is indicated in orange (resp. blue) next to the polar plot. If no proportion is indicated, then the proportions are 50/50. (**B**) Time series and polar plots of experiments in which the reference populations are synchronized (n=11). (**C**) Only one experiment underwent phase averaging.

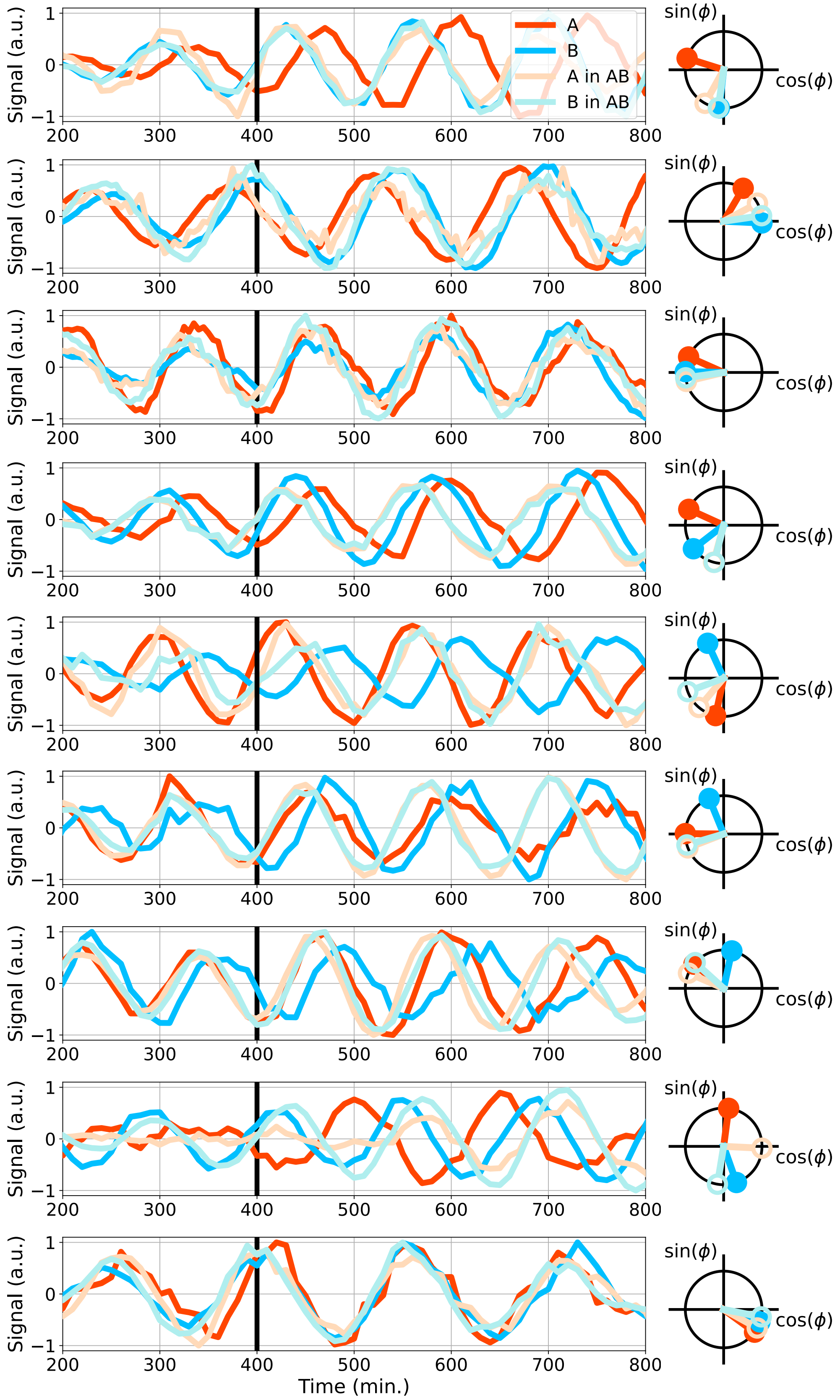


**Fig. S5.** Cells from both embryos are oscillating in the mixed population AB (n=9). Reference populations A and B are shown respectively in orange and blue. The oscillations of cells in the mixed population AB that came from the same embryo as reference population A (resp. B) are shown in light orange (resp. light blue). The thick black line indicates the time of the associated polar plot.

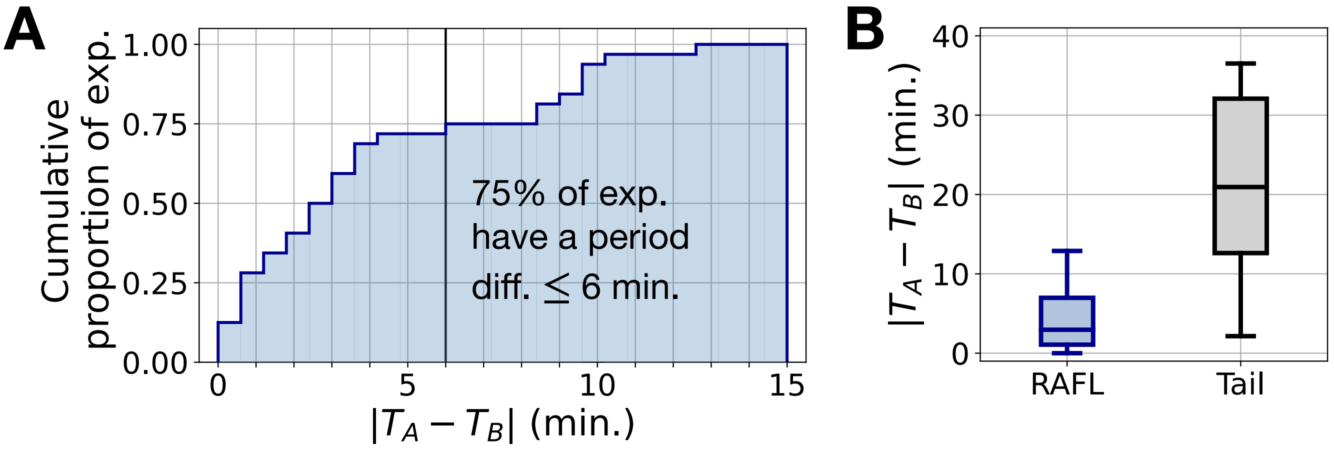


**Fig. S6.** Narrow period difference between populations A and B. (**A**) Cumulative proportion of pairs (n=32) of A and B populations as a function of the period difference between the two populations, in absolute value. 75% of experiments have a period difference $\leq$ 6 min. (**B**) The period difference distribution for pairs of A and B populations ("RAFL", blue, n=32) is much narrower than the distribution of period differences between the anterior PSM and the posterior PSM of the tail experiments described in Fig. S3 ("tail", grey, n=20).

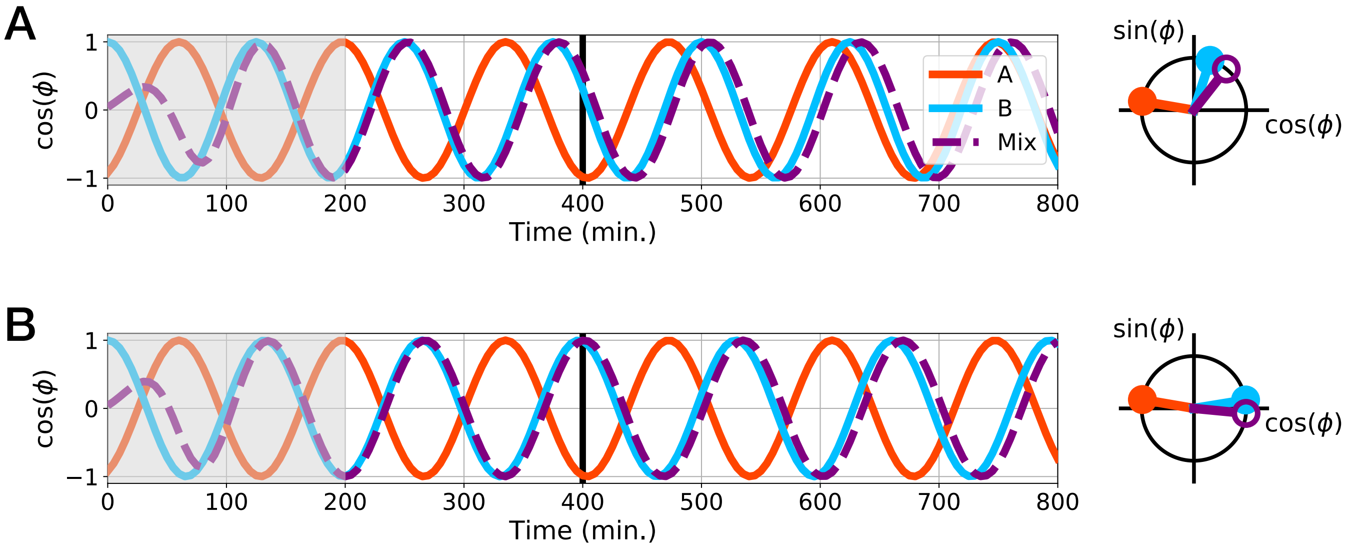


**Fig. S7.** Frequency differences can hide which oscillator was initially ahead. (**A-B**) Simulations of the rectified Kuramoto model with $\omega_B = 1.1\,\omega_A$ (**A**) and $\omega_B = 1.04\,\omega_A$ (**B**). Oscillators A and B are initially in anti-phase, with B slightly ahead of A. B wins, but appears behind A at $t = 400$ min. due to the frequency mismatch.

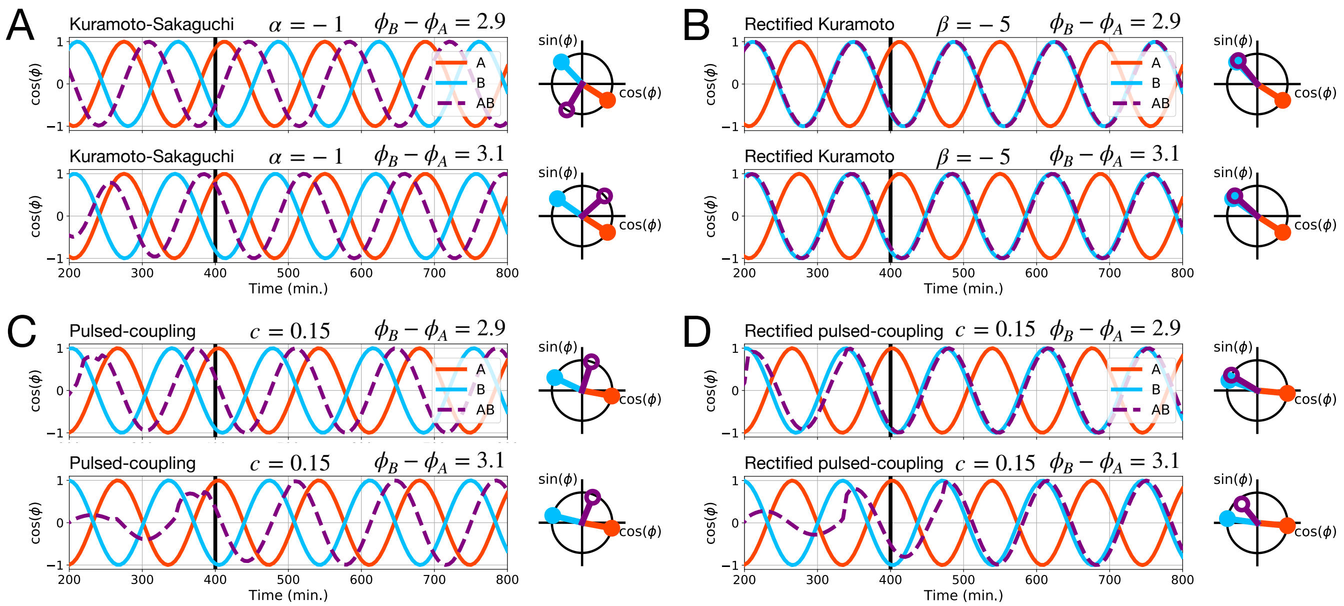


**Fig. S8.** Experiments with $|\phi_A - \phi_B| \approx \pi$ are the most informative to discriminate between models. (**A-D**) Different models' simulation outcomes for $|\phi_A - \phi_B| = 2.9$ and $3.1$. (**A**) Kuramoto-Sakaguchi model with $\alpha = -1$. (**B**) Rectified Kuramoto model with $\beta = -5$. (**C**) Pulsed-coupling model with strong coupling ($c = 0.15$). (**D**) Rectified pulsed-coupling model with strong coupling ($c = 0.15$).

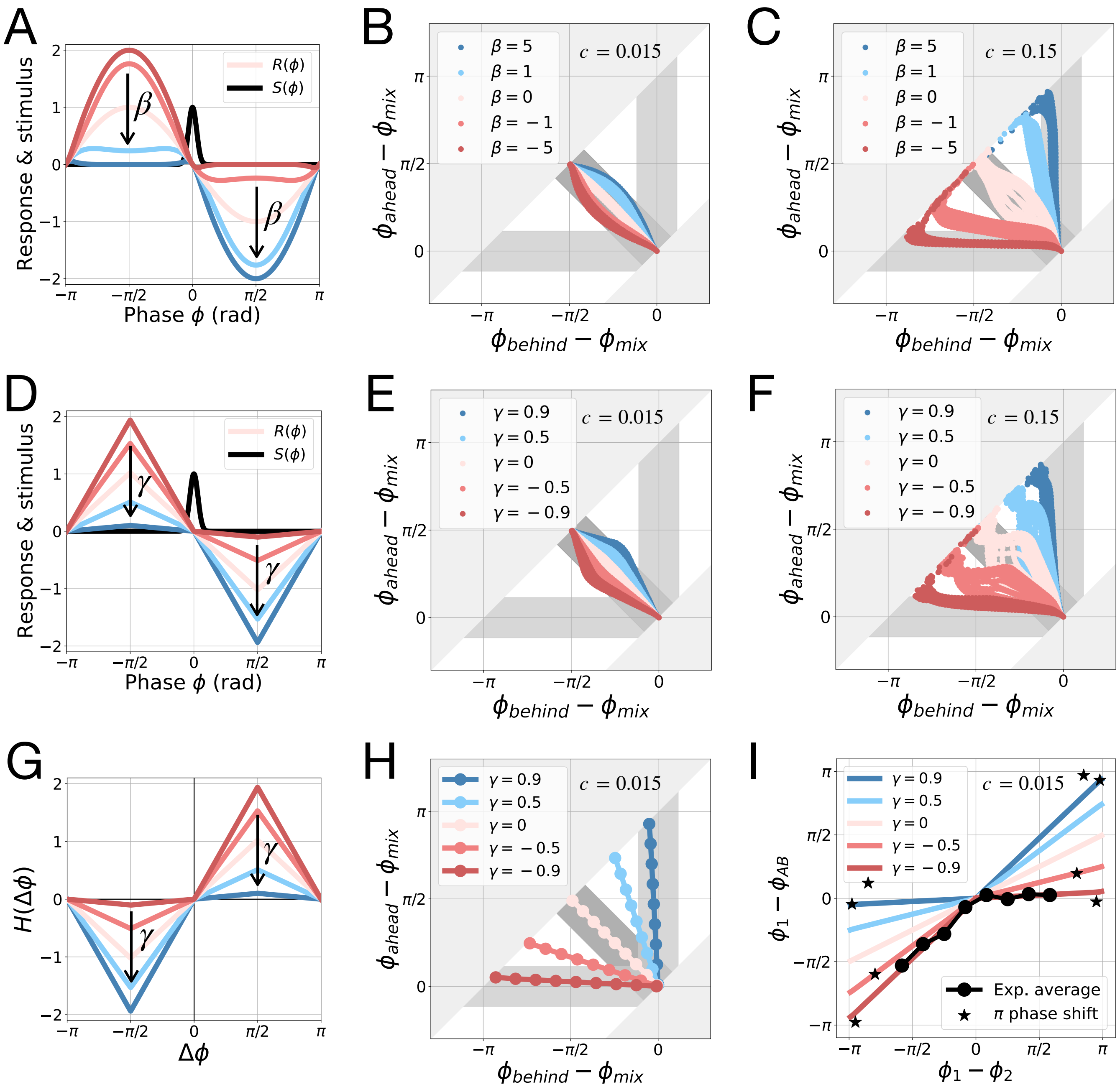


**Fig. S9.** Testing more alternative models. (**A-C**) Simulations of the rectified pulsed-coupling model. (**A**) Stimulus and rectified response function for different values of parameter $\beta$. (**B**) Simulation results of two-way synchronization with weak coupling strength. (**C**) Simulation results with strong coupling strength. (**D-F**) Simulations of the linear rectified pulsed-coupling model. (**D**) Stimulus and rectified response function for different values of parameter $\gamma$. (**E**) Simulation results of two-way synchronization with weak coupling strength. (**F**) Simulation results with strong coupling strength. (**G-I**) Simulations of the linear rectified Kuramoto model. (**G**) Coupling function for different values of parameter $\gamma$. (**H-I**) Simulation results of two-way synchronization with weak coupling strength, compared to experimental data (**I**). $\phi_1$ represents either $\phi_A$ or $\phi_B$. If $\phi_1 = \phi_A$, then $\phi_2 = \phi_B$, and vice versa. In experiments with a $\pi$ phase shift between $\phi_A$ and $\phi_B$, determining unambiguously which oscillator is ahead is impossible. For this reason, these experiments were not included in the data average.

**Table S1. Medium composition**

| Reagents | Culture medium | Dissection medium |
|---|---|---|
| DMEM/F-12 (Cell Culture Technologies | 50 mL | 50 mL) |
| BSA (Equitech-Bio, BAC62) | 0,02 g | 0,5 g |
| 1M HEPES (Gibco, 15630-106) | - | 850 µL |
| 10000 U/mL PenStrep (Gibco, 15140-122) | 50 µL | 850 µL |
| 45 % Glucose (Sigma, G8769) | 44.4 µL | 44.4 µL |
| 200mM L-Glutamine (Gibco, 25030-081) | 500 µL | 500 µL |

**Movie S1. Live fluorescence imaging of the embryonic oscillator ensemble shown in Fig. 2B.**

**Movie S2. Polar plots as a function of time for the four experiments shown in Fig. 3B.**
